# Subliminal Primes Bias the Spatial Locus of Involuntary Object Naming in a Two-Object Reflexive Imagery Task

**DOI:** 10.64898/2026.07.29.741423

**Authors:** Melisa Chulet, Jane Hamilda, Janu Rani, Kavitha Margabandhu, Thanusha K. Prasad, Christilda John Bosco, Sowmiya Sampathkumar, Mary Martine Victoria, Evelyn Sheena Sunderraj, Rasmi Rafi, Rohan Thomas Jepegnanam, George Abraham Ninan, Selvakumar Selvaganesan, Appaswamy Thirumal Prabhakar

## Abstract

Certain conscious contents—such as the covert name of a viewed object—arise involuntarily and resist suppression, a phenomenon captured by the Reflexive Imagery Task (RIT). Whether a subliminal prime can bias which of two simultaneously present objects captures such an involuntary naming response has not been established. We adapted a two-object RIT into a self-contained, browser-based instrument. Thirty-eight adults (M *age* = 21.6 years, SD = 2.3; 26 female) viewed 24 object pairs and were instructed to fixate a central cross, to refrain from thinking of the objects’ names, and to click an object whenever its name intruded into awareness. On each trial a masked prime (17 ms, flanked by pattern masks) was either the exact name of one object (Exact Word Prime), a semantic associate of one object (Semantic Prime), or a neutral string (No Prime), defining a primed side per trial. A post-experiment debriefing confirmed that participants noticed the masks but none consciously perceived or could identify the prime words, indicating that the primes were subliminal. Analyses excluded 91 of 912 trials (10.0%) with more than five clicks. The priming effect was robust across measures: an omnibus comparison of clicks across conditions was significant (Friedman χ²(2) = 6.59, p = .037); within both prime conditions participants clicked the primed side more than the not-primed side (Exact Word, p = .039; Semantic, p = .005); and the primed-to-not-primed ratio exceeded parity by roughly 57–59% (Laplace-smoothed ratio ≈ 1.58; one-sample Wilcoxon p < .001 for each). A Bayesian Poisson generalized linear mixed model with random participant intercepts confirmed a credible primed-side advantage (incidence rate ratio = 1.72, 95% credible interval [1.57, 1.90]) that did not differ between prime types. No general left/right response bias emerged. An exploratory right-side × Exact Word Prime interaction was inconsistent across model classes and is reported as hypothesis-generating. The findings indicate that subliminal lexical and semantic primes can steer the spatial locus of involuntary object naming.

## Introduction

A recurring puzzle in the science of consciousness is that many of the contents entering awareness are not deliberately summoned. Some conscious contents—images, urges, and subvocalizations—arise involuntarily and cannot be readily suppressed even when a person is explicitly instructed to keep them out of mind (Bargh & Morsella, 2008; Morsella, 2005; Morsella et al., 2016). The Reflexive Imagery Task (RIT; Allen et al., 2013) was developed to render this phenomenon experimentally tractable. In its canonical form, participants view a visual object and are instructed not to think of its name; despite this instruction, the object’s name is involuntarily subvocalized on roughly 80% of trials (Allen et al., 2013; Cho et al., 2018). The RIT effect is reliable and systematic, and it is sensitive to the properties of the eliciting stimulus and to the participant’s task set (Bhangal, Allen, et al., 2016; Bhangal, Cho, et al., 2016; Merrick et al., 2015).

Because the RIT externalizes an otherwise private event—the moment a name enters awareness—it provides a behavioral index of stimulus-elicited “involuntary entry” into consciousness (Baars, 1988; Dou et al., 2020). A natural extension asks whether the content that intrudes can be biased in advance, and in particular whether it can be biased by information the participant never consciously perceives. Priming offers a candidate mechanism: brief exposure to a word pre-activates its lexical and semantic representations and facilitates their subsequent processing, and this facilitation can occur even when the prime is masked and unavailable to report (Dehaene et al., 1998; Forster & Davis, 1984; Van den Bussche et al., 2009). Semantic priming, in which an associate rather than the item itself is presented, likewise pre-activates related representations through spreading activation, and meta-analytic evidence indicates that masked primes can be processed to a semantic level (Meyer & Schvaneveldt, 1971; Neely, 1991; Van den Bussche et al., 2009). If involuntary object naming behaves like other stimulus-driven cognition, a subliminal prime should increase the likelihood that the primed item, rather than a competing item, is the one whose name intrudes.

The two-object RIT variant is well suited to this question. When two objects are presented simultaneously to either side of fixation (Cho et al., 2018), the two lexical representations compete for entry into awareness, and the distribution of involuntary naming across the two sides becomes a sensitive measure of any bias in that competition. The present study embedded a masked-priming manipulation into such a task and verified, through post-experiment debriefing, that the primes were not consciously perceived. On each trial, a masked prime (presented for a single display frame between two pattern masks) was either the exact name of one of the two objects, a semantic associate of one object, or a neutral letter string. We asked three questions.

First, does a subliminal prime pull involuntary naming toward the primed object, relative to its competitor and to an unprimed baseline? Second, does an exact-word prime differ from a semantic prime in the magnitude of this pull? Third, independent of priming, is there any general tendency to respond more to one physical side than the other, and does the priming effect depend on which physical side happens to be primed?

Consistent with the ironic and reflexive character of the RIT (Wegner, 1994; Wegner et al., 1987), we predicted that instructing participants not to name the objects would nonetheless yield frequent intrusions, and that these intrusions would be drawn disproportionately toward the primed side. We report analyses in which extreme, perseverative-clicking trials were excluded, using distribution-free tests, population-averaged count models, and a mixed-effects model, to establish the robustness of the priming effect.

## Method

### Participants

Thirty-eight adults completed the task (M *age* = 21.6 years, SD = 2.3, range 18–29; one participant did not report age). Twenty-six participants identified as female and eleven as male (one did not report gender). Educational attainment ranged from high-school graduate to master’s degree, with the modal category being a bachelor’s degree (n = 24). Participants were recruited as a convenience sample and completed the task on their own web-enabled device. The study was conducted in accordance with the Declaration of Helsinki; all participants provided informed consent through the intake screen prior to testing. [Institutional ethics-committee approval number to be inserted.]

### Design

The experiment used a within-subjects design with a single three-level factor, Prime Condition (No Prime, Semantic Prime, Exact Word Prime). Each participant completed 24 recorded trials, 8 per condition, in a randomized order, yielding 912 trials across the sample. The primary dependent measures were the number of clicks (reported name intrusions) directed to each object and, on primed trials, the number of clicks to the primed versus the not-primed side.

### Materials and Apparatus

The task was implemented as a single, self-contained HTML/JavaScript instrument that ran in the participant’s web browser and logged all responses locally before export. Stimuli were drawn from a bank of 24 object pairs, each specifying a left object, a right object, and a semantic associate for each object (e.g., the pair apple–key was associated with fruit–lock; umbrella–candle with rain–flame). Whereas canonical RIT studies have typically used standardized line drawings (Snodgrass & Vanderwart, 1980), objects here were rendered as large, centrally legible emoji-style glyphs, one to the left and one to the right of a central fixation cross, against a uniform white field; administrative screens used a neutral, low-arousal visual style. Responses were made by clicking or touching an object, with touch and synthetic mouse events de-duplicated so that a single physical tap was logged once. Because the instrument was browser-based, exact stimulus timing depended on the participant’s display refresh rate and hardware, a limitation considered below.

### Procedure

After providing demographic information and consent, participants read the task instructions: to keep their eyes on the central fixation cross, to try not to think of the names of the two objects, and—if the name of an object nevertheless came to mind—to click or touch that object immediately, as many times as the name recurred. A single unrecorded warm-up trial preceded the 24 recorded trials.

Each trial followed a fixation → mask → prime → mask → stimulus sequence (Figure 1). A fixation cross was followed by a pattern mask (a 14-character random letter string, 100 ms), then the prime (17 ms, i.e., approximately one frame at a 60-Hz refresh rate), then a second pattern mask (100 ms). The two objects then appeared to the left and right of fixation for 6,000 ms, during which all clicks were recorded. In No Prime trials the primed slot contained a neutral string; in Exact Word Prime trials it contained the printed name of one of the two objects; in Semantic Prime trials it contained a semantic associate of one of the two objects. The primed side for each trial was defined as the side whose object corresponded to the prime word.

**Figure 1.**
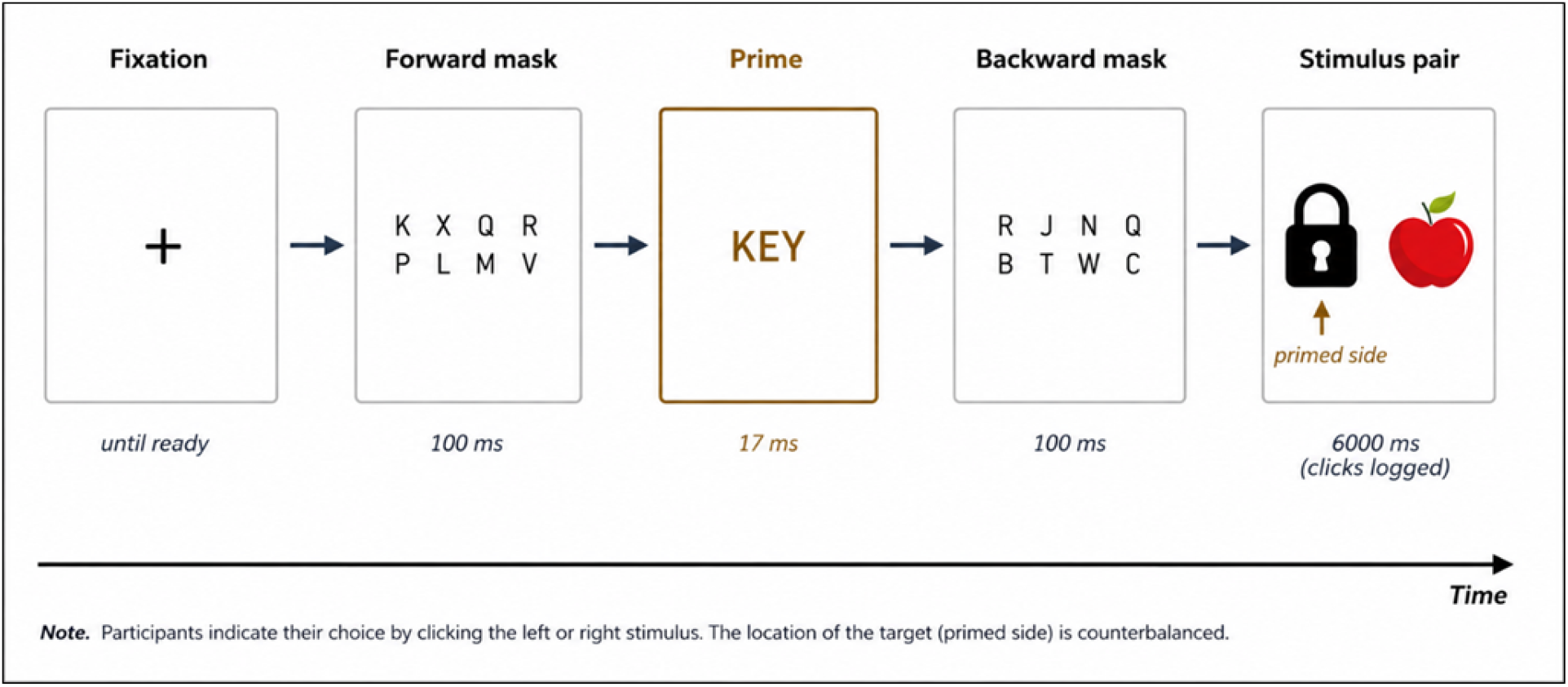
Schematic of the trial sequence. Each trial comprised a fixation cross, a forward pattern mask (100 ms), a masked prime presented for a single frame (17 ms), a backward pattern mask (100 ms), and the two-object stimulus display (6,000 ms), during which clicks to either object were logged. The prime word varied by condition (exact object name, semantic associate, or neutral string), defining the primed side on each primed trial.

### Prime-Awareness Debriefing (Manipulation Check)

Immediately after the task, each participant completed a structured debriefing interview probing awareness of the masked prime. Participants were asked whether they had noticed anything between the fixation cross and the appearance of the objects, whether they had seen any words, and whether they could report or guess any of the words presented. Every participant reported noticing the letter masks, but none reported consciously perceiving the prime words, and none could report or identify any prime. On this basis the prime presentations are treated as subliminal. This debriefing constituted as a subjective awareness measure.

### Data Reduction and Analysis

For each trial we recorded the number of clicks to the left and right objects, their sum (total clicks), the clicks directed to the primed and not-primed sides, and a Laplace-smoothed ratio of primed to not-primed clicks. Inspection of the raw data revealed a subset of extreme, perseverative-clicking trials in which a participant clicked one object many times (up to 36 clicks on a single trial); such trials dominated the untransformed means. We therefore excluded trials with more than five total clicks. This removed 91 of 912 trials (10.0%; No Prime = 23, Exact Word Prime = 34, Semantic Prime = 34), leaving 821 trials. After this exclusion, two participants had all eight of their trials in one condition removed and therefore lacked a valid mean in that cell; they were dropped from the complete-case omnibus test (leaving n = 36) but retained in all other analyses, which do not require a fully balanced participant × condition matrix.

Because click counts are non-negative, over-dispersed, and right-skewed, we combined distribution-free tests on participant-level means with count models on trial-level data. Omnibus differences across the three conditions were assessed with the Friedman test, followed by Wilcoxon signed-rank comparisons with Bonferroni correction. Within-condition primed-versus-not-primed differences were tested with Wilcoxon signed-rank tests, and the primed-to-not-primed ratio was tested against a value of 1 with a one-sample Wilcoxon test. To model trial-level counts, we fitted two complementary frameworks that account for the repeated-measures structure. First, population-averaged Poisson generalized estimating equations (GEE) with an exchangeable working correlation and robust standard errors, clustering on participant (Liang & Zeger, 1986). Second, subject-specific Poisson generalized linear mixed models (GLMMs) with a random intercept for participant, which explicitly partition between-participant variability from the fixed effects of interest (Baayen et al., 2008); these were estimated in a Bayesian framework by variational approximation, and we report posterior means, standard deviations, and 95% credible intervals (CrIs) for the fixed effects, together with incidence rate ratios (IRR = exp[b]). Because the variational approximation can underestimate posterior variance, GLMM credible intervals are interpreted alongside the GEE results. Analyses were conducted in Python using SciPy and statsmodels.

## Results

### Prime Awareness

The manipulation check confirmed the intended subliminality of the primes: all 38 participants reported awareness of the letter masks, whereas none reported seeing, and none could identify or guess, any of the prime words. All subsequent effects therefore reflect the influence of primes that were not consciously perceived.

### Descriptive Statistics

Participants responded on the majority of trials (M = 18.7 of 24 trials with at least one response). Mean first-response latencies were similar across conditions, ranging from 1,976 ms (Exact Word Prime) through 2,065 ms (Semantic Prime) to 2,209 ms (No Prime), consistent with responses that unfold over the 6-s viewing window rather than as immediate reactions. After excluding heavy-clicking trials, both prime types elicited more clicks to the (to-be) primed side than the per-side No Prime baseline (Table 1).

**Table 1.** Participant-Level Click Descriptives by Prime Condition (Filtered, ≤5-Click Trials)

| Condition | M | SD | Mdn | Max |
| --- | --- | --- | --- | --- |
| No Prime (baseline, per side) | 0.59 | 0.42 | 0.53 | 1.50 |
| Exact Word Prime (primed side) | 0.83 | 0.65 | 0.60 | 3.00 |
| Semantic Prime (primed side) | 0.85 | 0.82 | 0.63 | 4.00 |

*Note.* Values are means of participant-level means computed on trials with $\leq 5$ total clicks. For No Prime, the unit is mean clicks per side; for the prime conditions, the unit is mean clicks to the primed side. $n = 38$ participants.

### Effect of Prime Condition on Clicks to the Primed Side

The omnibus Friedman test comparing the three conditions was significant, χ²(2) = 6.59, p = .037 (n = 36 complete cases; Figure 2). Post-hoc Wilcoxon comparisons indicated that this reflected elevated clicking to the primed side under priming relative to the per-side baseline: No Prime versus Semantic Prime survived Bonferroni correction, W = 168.0, p = .048, and No Prime versus Exact Word Prime was of comparable magnitude but fell just short, W = 170.0, p = .052. The two prime types did not differ, W = 220.5, p = .845, and a Poisson GEE restricted to primed trials likewise found no difference between them in clicks to the primed side, b = −0.10, SE = 0.10, z = −0.96, p = .336.

**Figure 2.**
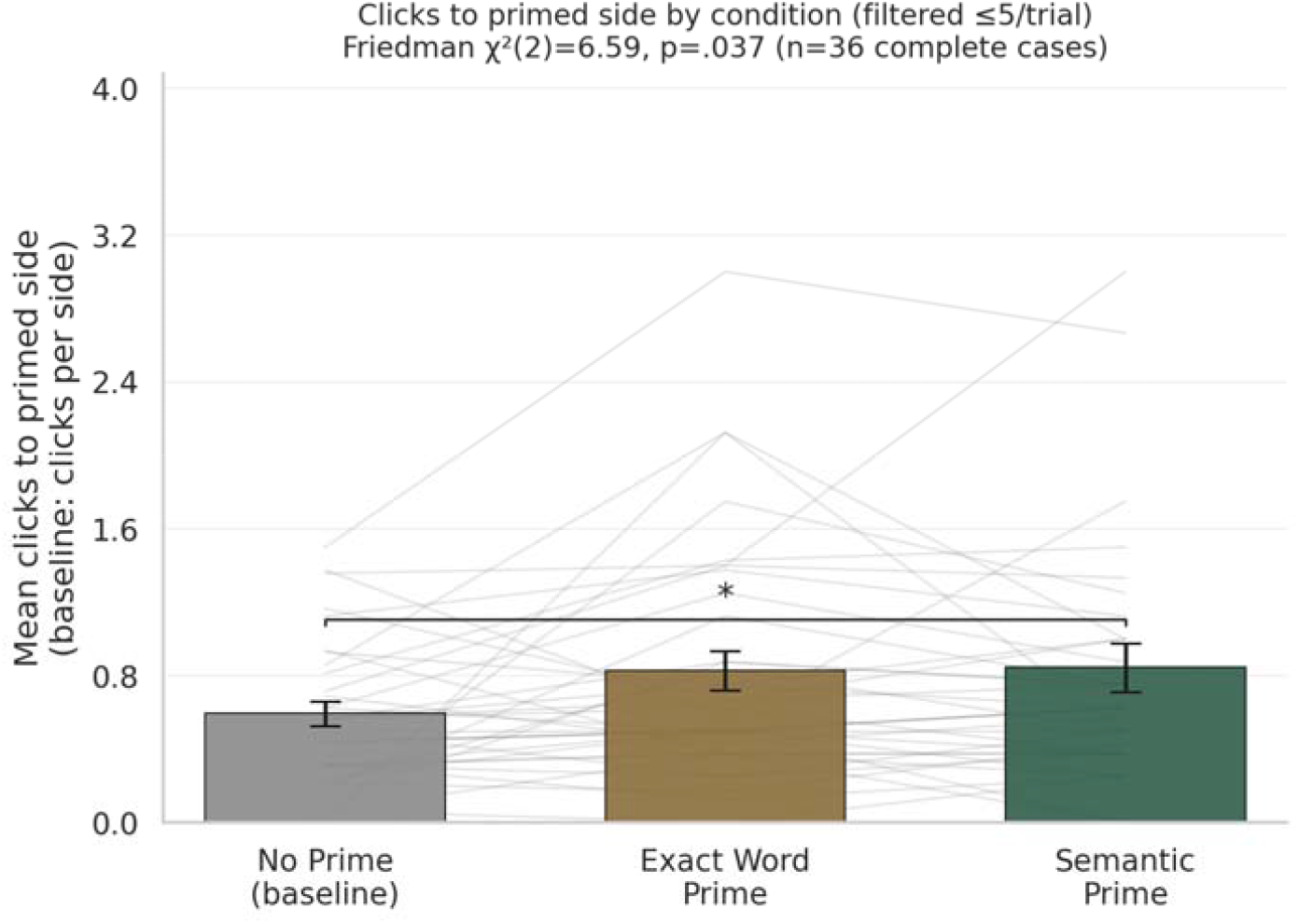
Mean clicks to the primed side (No Prime baseline expressed as mean clicks per side) by condition, filtered to ≤5-click trials. Grey lines link individual participants; error bars are ±1 SEM; the bracket marks the Bonferroni-significant No Prime versus Semantic Prime contrast. The three conditions differed overall (Friedman χ²[2] = 6.59, p = .037, n = 36 complete cases).

### Primed Versus Not-Primed Side

Within both prime conditions, participants directed reliably more clicks to the primed than to the not-primed side (Table 2). Under Exact Word Prime, M = 0.83 versus 0.51, W = 165.0, p = .039; under Semantic Prime, M = 0.85 versus 0.46, W = 132.5, p = .005. The corresponding primed-to-not-primed ratio departed strongly from parity in both conditions: participants clicked the primed side approximately 57–59% more often than the not-primed side (Exact Word Prime, M = 1.57; Semantic Prime, M = 1.59), and a one-sample Wilcoxon test rejected a ratio of 1 in each case (Ws = 19.0 and 38.0, both p < .001; Figure 4). Every measure—the omnibus test, the within-condition contrasts, and the ratio—therefore pointed to the same primed-side bias.

**Figure 3.**
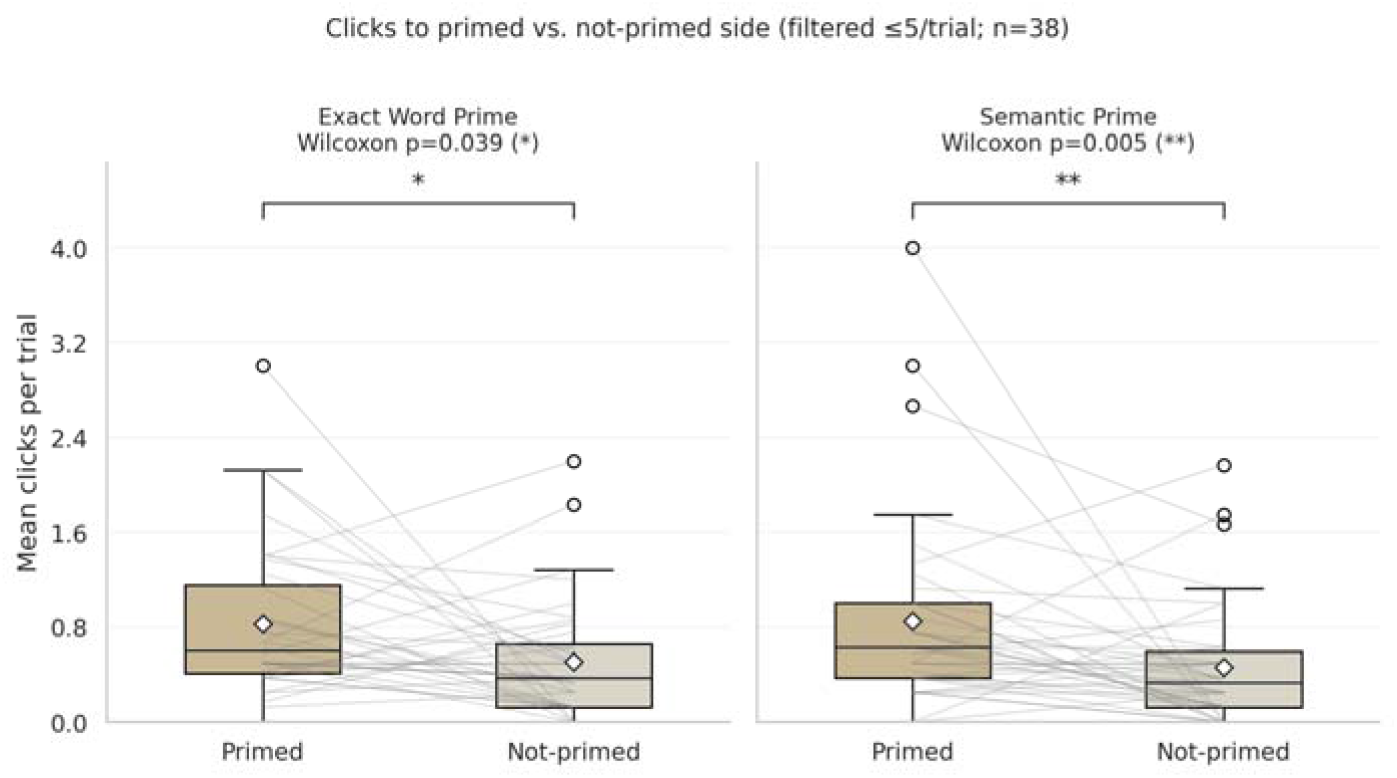
Clicks to the primed versus not-primed side within each prime condition (participant-level means, filtere ≤5-click trials, n = 38). Boxes show medians and interquartile ranges, diamonds show means, and grey lines lin individual participants. Both conditions show a reliable primed-side advantage (Exact Word, p = .039; Semantic, p = .005).

**Figure 4.**
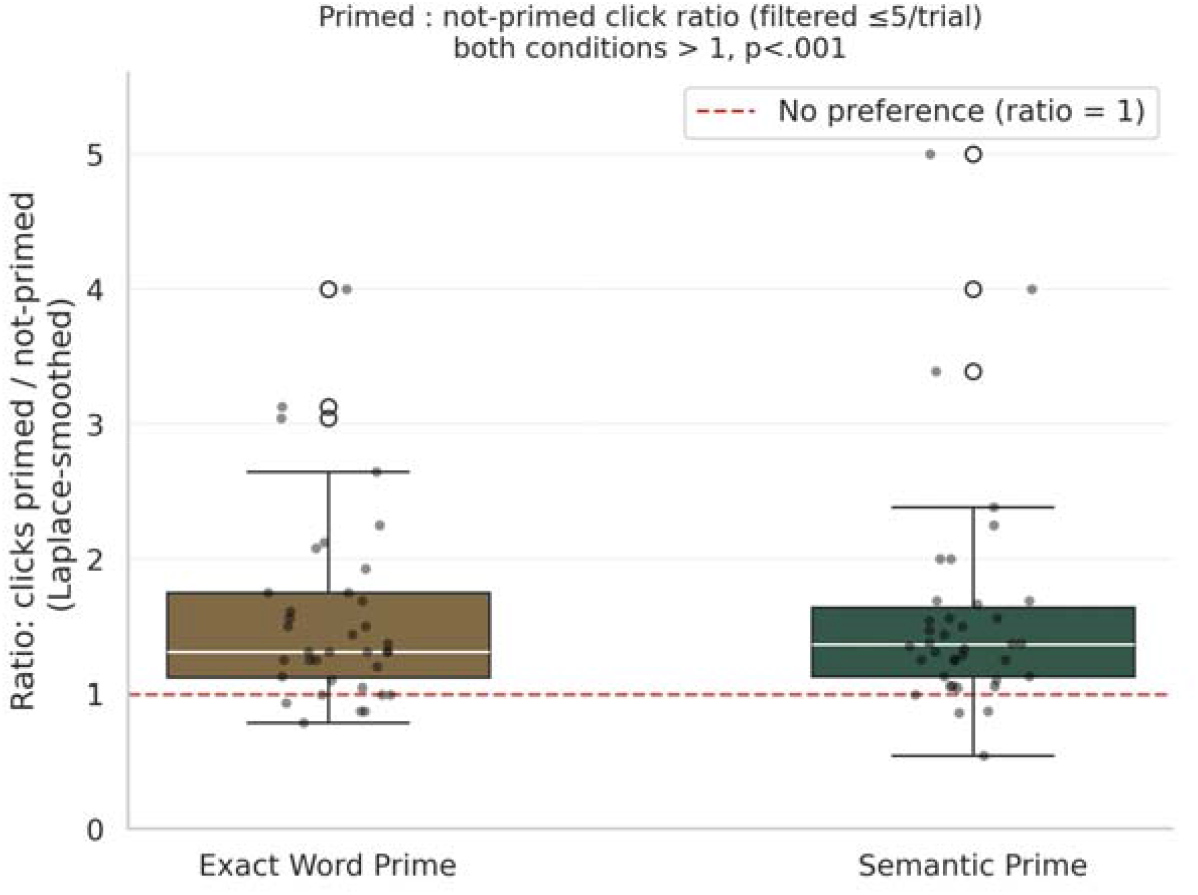
Primed-to-not-primed click ratio (Laplace-smoothed) by condition, filtered ≤5-click trials. The dashed line marks a ratio of 1 (no side preference). Both conditions differ significantly from 1 (one-sample Wilcoxon, p < .001).

**Table 2.** Within-Condition Primed Versus Not-Primed Comparisons and Ratio Tests (Filtered Data)

| Condition | M primed | M not-primed | W | p |
| --- | --- | --- | --- | --- |
| Exact Word Prime | 0.83 | 0.51 | 165.0 | **.039** |
| Semantic Prime | 0.85 | 0.46 | 132.5 | **.005** |
| *Ratio vs. 1: Exact Word* | *1.57* | — | 19.0 | **< .001** |
| *Ratio vs. 1: Semantic* | *1.59* | — | 38.0 | **< .001** |

Wilcoxon signed-rank tests on participant-level means ( $n = 38$ per condition). “Ratio” rows report the mean Laplace-smoothed primed-to-not-primed ratio and a one-sample Wilcoxon test against a value of 1.

### Mixed-Effects Model of the Primed-Side Advantage

To model the primed-side effect at the trial level while accommodating between-participant differences in overall click rate, we fitted a Poisson GLMM to the filtered primed trials in long format (each trial contributing a primed-side and a not-primed-side count), with side and prime condition and their interaction as fixed effects and a random intercept for participant (1,080 observations, 38 participants). The random participant intercept was sizeable (SD = 0.57 on the log scale), confirming that participants differed markedly in baseline click rate and justifying the mixed-model approach. Controlling for this heterogeneity, clicks to the primed side were credibly elevated relative to the not-primed side, b = 0.54, 95% CrI [0.45, 0.64], IRR = 1.72 [1.57, 1.90] (Table 3). This primed-side advantage did not differ between the two prime types (side × condition interaction, b = −0.12, 95% CrI [−0.26, 0.02]), nor was there a credible main effect of prime type (b = 0.01, 95% CrI [−0.10, 0.12]). The mixed model thus converges with the population-averaged GEE, in which the same primed-side contrast was also significant, b = 0.54, SE = 0.20, z = 2.69, p = .007.

**Table 3.** Bayesian Poisson GLMM of Clicks to the Primed Versus Not-Primed Side (Filtered Data)

| Fixed effect | b | SD | 95% CrI | IRR |
| --- | --- | --- | --- | --- |
| Intercept (not-primed, Exact Word) | -0.86 | 0.04 | [-0.94, -0.79] | 0.42 |
| **Side (primed vs. not-primed)** | **0.54** | 0.05 | **[0.45, 0.64]** | **1.72** |
| Prime condition (Semantic vs. Exact) | 0.01 | 0.06 | [-0.10, 0.12] | 1.01 |
| Side × Semantic Prime | -0.12 | 0.07 | [-0.26, 0.02] | 0.89 |

Poisson GLMM with a random intercept for participant (random-intercept SD = 0.57 on the log scale), estimated by variational Bayes. $b$ = posterior mean log-rate coefficient; CrI = credible interval; IRR = incidence rate ratio ( $\exp[b]$ ). Bold row: the primed-side effect, whose CrI excludes zero. $n = 1,080$ side-level observations from 38 participants (primed trials, $\leq 5$ clicks).

### General Left/Right Bias and Physical-Side Effects

A separate set of analyses asked whether, independent of priming, participants favored one physical side. No general left/right response bias emerged, either across all trials (M *left* = 0.62, M *right* = 0.66; W = 262.5, p = .179) or within any single condition (all p > .08; Table 4, Figure 5). A Poisson GEE modeling clicks as physical side (left vs. right) crossed with prime condition yielded a right-side × Exact Word Prime interaction that was only a non-significant trend, b = 0.25, SE = 0.14, z = 1.76, p = .079, with no comparable effect for the Semantic Prime (p = .999); a parallel Poisson GLMM returned a credible interaction of similar sign (b = 0.25, 95% CrI [0.11, 0.39], IRR = 1.29). Because this term reached credibility in the mixed model but not significance in the population-averaged model, and was not corroborated by a direct test of whether the priming pull depended on the primed side’s physical location (b = 0.20, p = .116; interaction p = .547), we regard it as inconsistent and hypothesis-generating rather than an established effect.

**Table 4.**
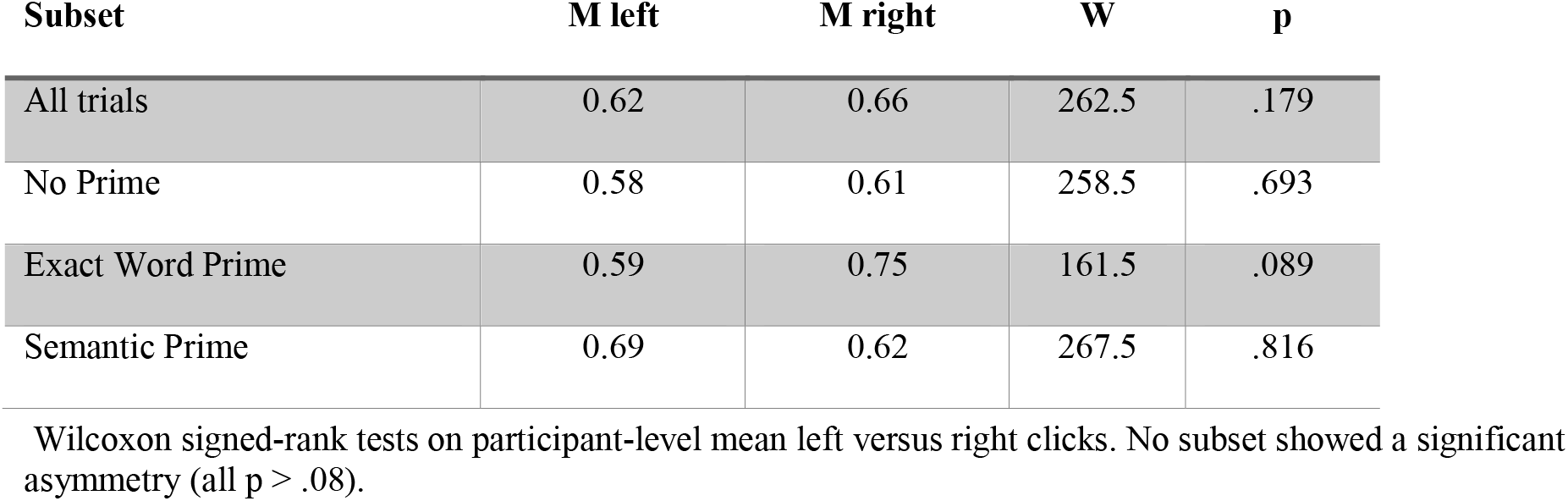
General Left Versus Right Click Bias by Subset (Filtered Data)

| Subset | M left | M right | W | p |
| --- | --- | --- | --- | --- |
| All trials | 0.62 | 0.66 | 262.5 | .179 |
| No Prime | 0.58 | 0.61 | 258.5 | .693 |
| Exact Word Prime | 0.59 | 0.75 | 161.5 | .089 |
| Semantic Prime | 0.69 | 0.62 | 267.5 | .816 |

Wilcoxon signed-rank tests on participant-level mean left versus right clicks. No subset showed a significant asymmetry (all $p > .08$ ).

**Figure 5.**
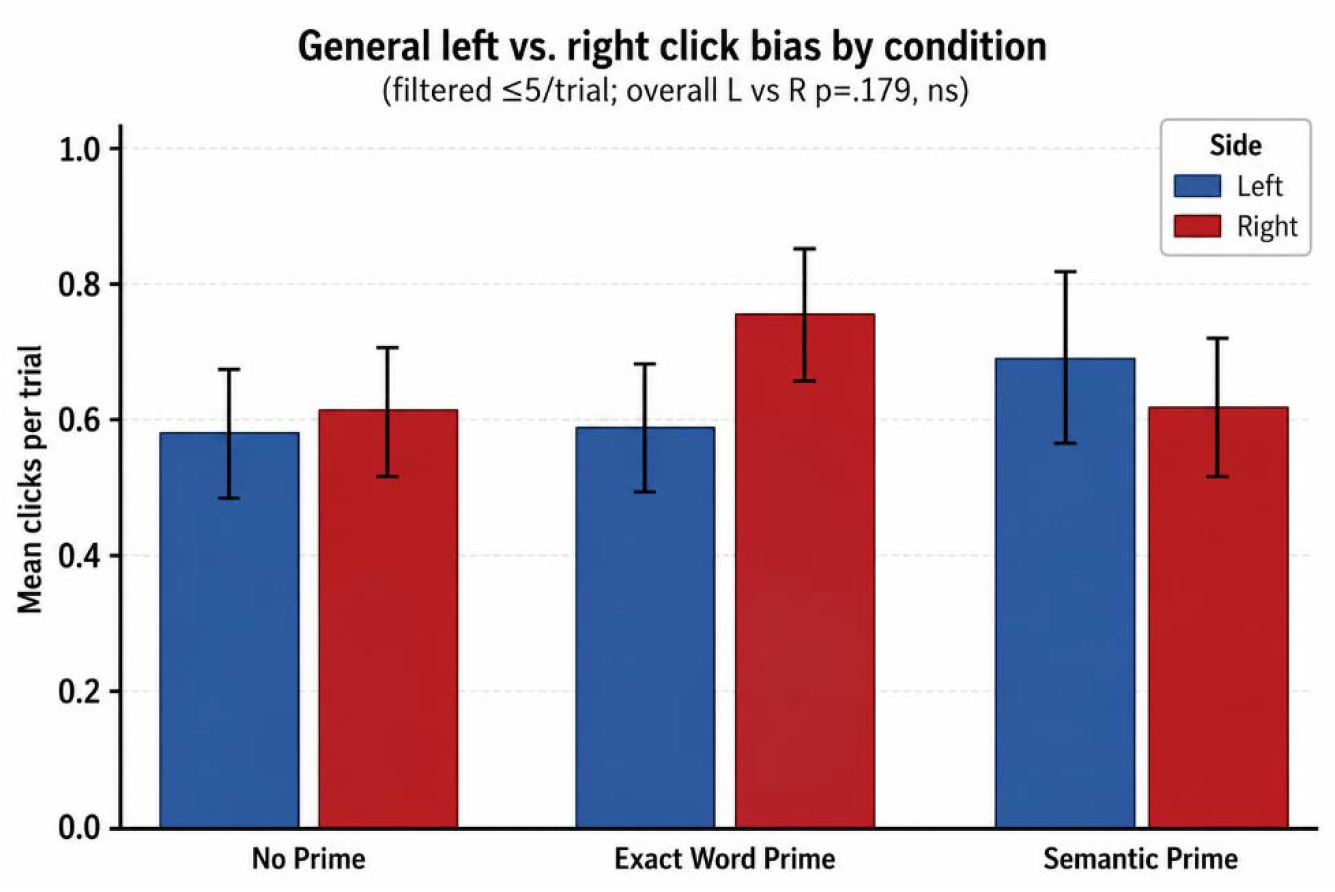
Mean clicks to the left versus right side by prime condition (participant-level means, filtered data; error bars ±1 SEM). No condition shows a significant left/right asymmetry on its own; the overall comparison was non-significant (p = .179).

**Figure 6.**
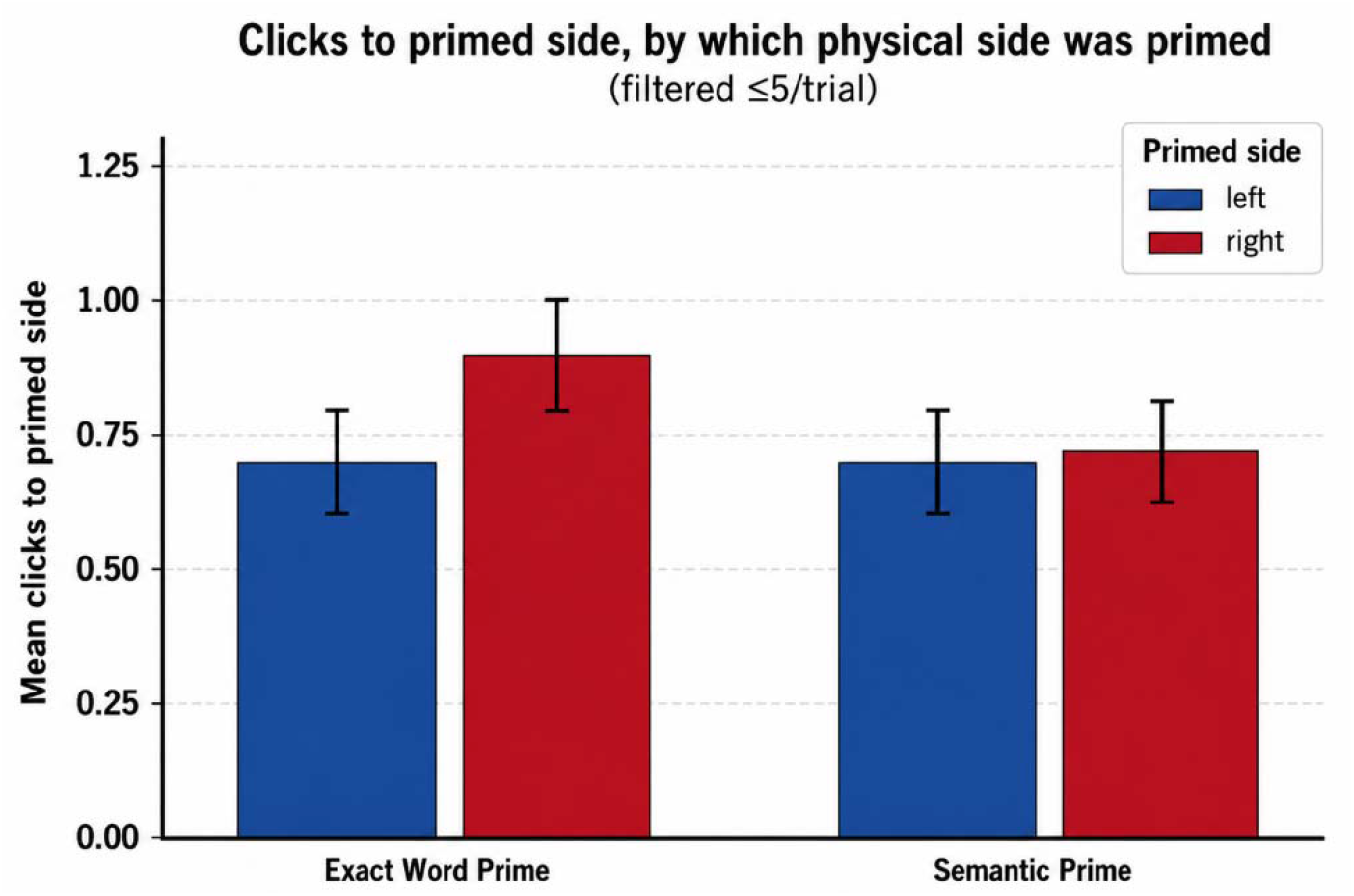
Mean clicks to the primed side split by whether the primed side happened to be physically left or right (filtered data; error bars = SEM). The numerical right-over-left difference did not reach significance (p = .116) and did not differ between prime types (interaction p = .547).

## Discussion

Our study demonstrates that involuntary object naming is not determined solely by consciously available visual information, but can be selectively biased by information that remains outside conscious awareness. Using a masked-prime version of the Reflexive Imagery Task, we found that subliminal primes systematically shifted the spatial locus of involuntary naming toward the primed object, with comparable effects following both exact-word and semantic primes. The convergence of these effects across complementary analyses provides evidence that subliminal information can influence which of competing object representations gains access to involuntary conscious content.

In this larger sample the effect was robust across every analytic approach: the omnibus comparison of clicks across conditions was significant, both within-condition primed-versus-not-primed contrasts were significant, the primed-to-not-primed ratio departed from parity at p < .001 in each condition, and a mixed model with random participant intercepts returned a credible primed-side advantage (IRR = 1.72). Because the debriefing established that the primes were subliminal, the central result is that a subliminal prime can steer the spatial locus of involuntary object naming toward the primed item.

The finding extends the RIT literature in two ways. First, it shows that the content that involuntarily enters awareness in the RIT is not fixed by the visible stimuli alone but can be biased by a preceding, unseen prime, complementing prior demonstrations that the RIT effect is shaped by stimulus salience, set, and learning history (Bhangal, Allen, et al., 2016; Bhangal, Cho, et al., 2016; Merrick et al., 2015). Second, the equivalence of exact-word and semantic primes—neither the post-hoc contrasts, the GEE, nor the GLMM interaction distinguished them—suggests that the bias operates at least partly at a semantic rather than a purely lexical level. This is consistent with spreading-activation accounts of semantic priming (Meyer & Schvaneveldt, 1971; Neely, 1991) and with meta-analytic evidence that masked primes can be processed semantically (Dehaene et al., 1998; Van den Bussche et al., 2009). That a subliminal semantic associate was as effective as the object’s own name in biasing which name intruded is a non-trivial constraint on the mechanism.

The convergence across measures is itself informative. In a smaller pilot sample, the effect had been detectable mainly through within-participant contrasts—the ratio and the mixed model—while the omnibus comparison of absolute counts was under-powered. With 38 participants the primed-side bias is now evident even in the omnibus test and the raw within-condition comparisons, indicating that the earlier null omnibus reflected limited power rather than the absence of an effect. The sizeable random participant intercept (SD = 0.57) confirms substantial individual differences in overall click rate, so relative, within-participant contrasts and mixed-effects models remain the most efficient way to characterize the effect.

The side analyses were largely reassuring: there was no general left/right response bias in any condition, indicating that the primed-side effect reflects the priming manipulation rather than a spatial-motor habit. A right-side × Exact Word Prime interaction reached credibility in the mixed model but was only a trend in the population-averaged model and was not corroborated by a direct test; we therefore do not interpret it as an established effect, and flag it for targeted replication.

### Limitations

Several limitations remain. Although the sample is now moderate (N = 38), it is homogeneous (young adults), which constrains generalizability. The task was delivered through participants’ own browsers, so the nominal 17-ms prime duration could not be guaranteed across devices with differing refresh rates and rendering pipelines; effective prime exposure likely varied between participants. Although the debriefing indicated that no participant consciously perceived the primes, this is a subjective, retrospective awareness measure; it does not have the sensitivity of an objective, trial-level visibility test (e.g., a forced-choice prime-identification task yielding d′), which provides the most rigorous evidence for subliminality (Merikle et al., 2001; Van den Bussche et al., 2009). We therefore describe the primes as subliminal by subjective report while noting that minimal residual visibility on some trials cannot be fully excluded. The dependent measure relies on participants’ introspective reports of fleeting conscious contents, which are subject to known inaccuracies and demand characteristics (Nisbett & Wilson, 1977; Wegner, 1994). Finally, perseverative clicking on a minority of trials required a principled but post hoc exclusion rule, and the several exploratory side × condition contrasts raise the risk of false positives.

### Future Directions and Conclusion

Future work should include an objective prime-visibility check (forced-choice identification with d′), tighter control of stimulus timing, and a more diverse sample, and could pair the paradigm with electrophysiology to time-lock the involuntary entry of the primed name (Dou et al., 2020). Systematically varying prime–target semantic distance would help localize the level at which the bias operates. In sum, within a two-object reflexive imagery task, subliminal exact-word and semantic primes reliably biased which object’s name involuntarily entered awareness—an effect now convergent across omnibus, within-condition, ratio, and mixed-effects analyses—reinforcing the view that even seemingly self-generated conscious contents are substantially subject to external, stimulus-based control.

## Declarations

### Ethics approval

Approved by the Institutional Review Board and Ethics Committee of Christian Medical College, Vellore (IRB Min. No. 2508114, dated 20.08.2025). All procedures accorded with the Declaration of Helsinki, and written informed consent was obtained from all participants or their surrogates.

### Data and code availability

Data and code used in this analysis is freely available at https://osf.io/vrb7f

### CRediT Authorship Contribution Statement

**Melisa Chulet:** Conceptualization, Writing – review & editing. **Jane Hamilda:** Investigation, Writing – review & editing. **Janu Rani:** Investigation, Writing – review & editing. **Kavitha Margabandhu:** Software, Writing – review & editing. **Thanusha K. Prasad:** Investigation, Writing – review & editing. **Christilda John Bosco:** Investigation, Writing – review & editing. **Sowmiya Sampathkumar:** Investigation, Writing – review & editing. **Mary Martine Victoria:** Investigation, Writing – review & editing. **Evelyn Sheena Sunderraj:** Investigation, Writing – review & editing. **Rasmi Rafi:** Investigation, Writing – review & editing. **Rohan Thomas Jepegnanam:** Software, Writing – review & editing. **George Abraham Ninan:** Investigation, Writing – review & editing. **Selvakumar Selvaganesan:** Investigation, Writing – review & editing. **Appaswamy Thirumal Prabhakar:** Formal analysis, Writing – original draft, Writing – review & editing.

